# Early mannitol-triggered changes in the Arabidopsis leaf (phospho)proteome

**DOI:** 10.1101/264259

**Authors:** Natalia Nikonorova, Lisa Van den Broeck, Shanshuo Zhu, Brigitte van de Cotte, Marieke Dubois, Kris Gevaert, Dirk Inzé, Ive De Smet

**Author notes:** CORRESPONDING AUTHOR: Ive De Smet, VIB-UGent Center for Plant Systems Biology, Technologiepark 927, 9052 Gent, Belgium. Equal contribution (Natalia Nikonorova, Lisa Van den Broeck). Present address: Institut de Biologie Moléculaire des Plantes, CNRS, 67084 Strasbourg, France.

## Abstract

Drought is one of the most detrimental environmental stresses to which plants are exposed. Especially mild drought is relevant to agriculture and significantly affects plant growth and development. In plant research, mannitol is often used to mimic drought stress and study the underlying responses. In growing leaf tissue of plants exposed to mannitol-induced stress, a highly-interconnected gene regulatory network is induced. However, early signaling and associated protein phosphorylation events that likely precede part of these transcriptional changes are largely unknown. Here, we performed a full proteome and phosphoproteome analysis on growing leaf tissue of *Arabidopsis* plants exposed to mild mannitol-induced stress and captured the fast (within the first half hour) events associated with this stress. Based on this in-depth data analysis, 167 and 172 differentially regulated proteins and phosphorylated sites were found back, respectively. Additionally, we identified H(+)-ATPASE 2 (AHA2) and CYSTEINE-RICH REPEAT SECRETORY PROTEIN 38 (CRRSP38) as novel regulators of shoot growth under osmotic stress.

**Highlight:** We captured early changes in the *Arabidopsis* leaf proteome and phosphoproteome upon mild mannitol stress and identified AHA2 and CRRSP38 as novel regulators of shoot growth under osmotic stress

## INTRODUCTION

Plants are exposed to a range of disadvantageous environmental conditions, which has led to the development of various molecular coping mechanisms (Wang *et al.*, 2017*a*; Berens *et al.*, 2017; Demarsy *et al.*, 2017). Drought is one of the most detrimental environmental stresses to which plants are exposed (Boyer, 1982; Wang *et al.*, 2003). Plant growth and subsequently plant yield are drastically decreased as a result of drought. In a temperate climate, water scarcity rarely threatens the survival of the plant, but rather reduces the growth and yield of the crop. Undoubtedly, the underlying molecular mechanisms differ depending on how severe the water limitation is, as plant lines that are more tolerant to severe stress rarely perform better under mild stress (Skirycz *et al.*, 2011*c*). Because mild drought is more relevant to agriculture and significantly affects plant growth and development, studies on growth responses of plants exposed to mild drought are of increasing importance (Aguirrezabal *et al.*, 2006; Pereyra-Irujo *et al.*, 2008; Clauw *et al.*, 2015). Several studies have focussed on unravelling the transcriptomic changes associated with the drought response (Harb *et al.*, 2010; Clauw *et al.*, 2015, 2016; Bac-Molenaar *et al.*, 2016; Rasheed *et al.*, 2016; Dubois *et al.*, 2017; Verslues, 2017) and a few studies even addressed drought response on a proteome or phosphoproteome level (Singh and Jwa, 2013; Katam *et al.*, 2016; Vu *et al.*, 2016).

Capturing the early response upon drought, a condition that builds up gradually, is not straightforward. Because drought is associated with osmotic stress, *in vitro* alternatives, such as mannitol, sorbitol or polyethylene glycol (PEG), are used to study the molecular events associated with this stress condition (Verslues *et al.*, 2006*a*). By transferring plants at a desired time point during development to an osmotic compound or by adding such a compound to liquid cultures, the very early signalling mechanisms associated with osmotic stress can be revealed. Low concentrations of mannitol (25 mM) induce mild stress, triggering a decrease in Arabidopsis rosette size of approximately 50% without affecting the plant’s development or survival (Claeys *et al.*, 2014). Mannitol can therefore be used as a growth-repressive compound and has been shown to be ideal for studying growth-regulating events (Skirycz *et al.*, 2011*c*; Van den Broeck *et al.*, 2017).

The osmotic stress responses in young growing leaves are very different from those in mature leaves. For example, in young leaves, mild stress induces the rapid accumulation of the ethylene precursor ACC (1-aminocyclopropane-1-carboxylate) and presumably also ethylene itself, instead of the classic drought-related hormone abscisic acid (ABA) (Skirycz *et al.*, 2011*a*). Unravelling the growth regulation upon stress is thus preferably studied in growing tissue instead of mature leaves or whole seedlings, as growing tissues are more subdued by growth inhibitory mechanisms. In growing leaf tissue exposed to mannitol induced stress, a highly-interconnected gene regulatory network (GRN) is induced. The transcription factors that are part of this network regulate each other’s expression and have been shown to regulate leaf growth upon osmotic stress (Van den Broeck *et al.*, 2017). Some members of this GRN, such as ETHYLENE RESPONSE FACTOR 6 (ERF6), ERF9 and WRKY15, can activate the expression of *GIBBERELLIN2-OXIDASE6* (*GA2-OX6*), a gene encoding a gibberellin degradation enzyme (Rieu *et al.*, 2008; Dubois *et al.*, 2013; Van den Broeck *et al.*, 2017). This results in decreased levels of gibberellin and DELLA protein stabilisation, which pushes the cells to permanently exit the cell division phase and into the cell differentiation phase (Claeys *et al.*, 2012). The first transcriptional changes occur very rapidly, after 40 minutes of stress. However, early signalling and associated phosphorylation events that precede part of these transcriptional changes are largely unknown, likely because most phosphoproteomic studies focused on severe, lethal stress or on whole seedlings, masking the growth-specific phosphorylation events (Bhaskara *et al.*, 2017*a*).

While the transcriptional events orchestrating leaf growth upon mild stress have been studied extensively, the early proteome and phosphoproteome changes are not yet fully understood. In this study, we performed full proteome and phosphoproteome analyses on growing leaf tissue exposed to mannitol-induced stress and captured the rapid (within the first half hour) events associated with this stress. We demonstrate differences in proteome changes in the early (30 min) and later (4 h) mannitol-triggered proteome, such as the translational machinery and oxidation-reduction processes. Next, we evaluated the phoshoproteome and found several connections with the GA¯DELLA pathway, which form interesting candidates for follow-up studies. We compared four phosphoproteome datasets after mild and severe stress highlighting their distinct signalling pathways. Finally, through validation of some candidates for a growth phenotype upon mild mannitol treatment, we identified H(+)-ATPASE 2 (AHA2) and CYSTEINE-RICH REPEAT SECRETORY PROTEIN 38 (CRRP38) as novel regulators of osmotic stress signalling and response.

## MATERIAL AND METHODS

### Plant material and growth conditions

Wild-type plants (Col-0) were grown *in vitro* at 21°C under a 16-h-day (110 mmol/(m^2^s)) and 8-h-night regime. 64 wild-type seeds were sown on a 14-cm-diameter Petri dish with solid ½ MS medium (Murashige and Skoog, 1962; 6.5 g/L agar, Sigma), overlaid with a nylon mesh (Prosep) of 20-μm pore size. During growth, plates were randomized. For the short-term (30 min mannitol stress) proteome and phosphoproteome analyses, half of the plants were transferred to ½ MS medium containing 25 mM mannitol (Sigma) at 15 days after stratification (DAS). The third leaf was harvested after 30 min. The other half of the plants was not transferred, and the third leaf was harvested before transfer (time point 0). In total, 4 biological repeats were performed. For the proteomic experiment after 4 h of mannitol stress and the expression analysis, half of the plants were transferred to control ½ MS medium, the other half to ½ MS medium containing 25 mM mannitol at 15 DAS. The third leaf was harvested 20 min and 40 min (for expression analysis) or 4 h (for proteomics) after transfer. In total 3 biological replicates were performed, and approximately 100 mg leaf material was harvested per sample. All experiments were performed independently.

### qPCR analyses

Samples were immediately frozen in liquid nitrogen and ground with a Retsch machine and 3-mm metal beads. Subsequently, RNA was extracted with TriZol (Invitrogen) and further purified with the RNeasy plant mini kit (Qiagen). For cDNA synthesis, the iScript cDNASynthesis Kit (Bio-Rad) was used with 1 µg of RNA as starting material. qRT-PCR was performed with the LightCycler 480 Real-Time SYBR Green PCR System (Roche). The data were normalised against the average of housekeeping genes AT1G13320 and AT2G28390 (Czechowski, 2005), as follows: dCt = Ct (gene) – Ct (average [housekeeping genes]) and ddCt = dCt (Control) – dCt (Treatment). Ct represents the number of cycles at which the SYBR Green fluorescence reached a threshold during the exponential phase of amplification. Primers were designed with Primer-BLAST (https://www.ncbi.nlm.nih.gov/tools/primer-blast/) (**Supplementary Information**).

### (Phospho)proteome workflow

Plant material was flash-frozen in liquid nitrogen and ground into a fine powder. Subsequent proteome and phosphoproteome analyses were performed as previously described (Vu *et al.*, 2016; Nikonorova *et al.*, 2018). For details, we refer to the **Supplementary Information**. LC-MS/MS analysis was performed as previously described (Vu *et al.*, 2016). Both the proteome and phosphoproteome samples were analysed using 3 h gradients on a quadrupole Orbitrap instrument (Q Exactive).

MS/MS spectra were searched against the Arabidopsis proteome database (TAIR10, containing 35,386 entries; http://www.arabidopsis.org/) using the MaxQuant software (version 1.5.4.1). Details on settings can be found in **Supplementary Information**. All MS proteomics data have been deposited to the ProteomeXchange Consortium via the PRIDE partner repository with the data set identifier PXD008900. For the quantitative proteome and phosphoproteome analyses, the “ProteinGroups” and ‘Phospho(STY)sites’ output files, respectively, generated by the MaxQuant search were loaded into Perseus software. For phosphoproteome data only high-confidence hits with phosphorylation localisation probabillity > 0.75 were included in analysis. Data analysis was performed as described previously (Vu *et al.*, 2016), and modifications are added in the main text.

### In silico analyses, data visualisation and statistics

To generate networks for known and predicted protein-protein interactions, the datasets were loaded to the STRING database (https://string-db.org; version 10.5) using high confidence interaction score (> 0.7). As active interaction sources text mining, experiments, databases, co-expression, neighbourhood, gene fusion and co-occurrence were selected. (Phospho)proteins that did not have any interaction were removed from the network. Biological Process terms were retrieved from the TAIR portal (www.arabidopsis.org, Bulk Data Retrieval). Further data visualisation was performed in Cytoscape (version 3.5.1) on the extracted interaction network and GO annotations. GO-enrichment analysis was performed in PLAZA 4.0 workbench using entire species as background model and 0.05 as a p-value cut-off. The HMMER web server was used for the prediction of kinase domains. For Motif-X analyses (Chou and Schwartz, 2011), the sequences (limited to 13 amino acids) of up- and downregulated phosphosites were pre-aligned with the phosphosite centered. The IPI Arabidopsis Proteome was used as the background database. The occurrence threshold was set at the minimum of 5 peptides, and the p-value threshold was set at <10^−6^. Venn diagrams were created using the Venny 2.1 online tool (http://bioinfogp.cnb.csic.es/tools/venny). Barcharts, boxplots and statistical analyses for qPCR and phenotyping data were performed using R software (https://www.R-project.org) and all figures were generated with Inkscape or Photoshop. Details on ANOVA results can be found in **Supplementary Information**.

### Genotyping

The *crrsp38-1* (SALK_151902) and *aha2-4* (SALK_082786) mutant *Arabidopsis* plants were obtained from Nottingham Arabidopsis Stock Centre (NASC). The homozygous *Arabidopsis* mutants were identified by PCR using primers specific to the insertion T-DNA (**Supplementary Information**) and the LB primer (ATTTTGCCGATTTCGGAAC). The SALK lines used are in Col-0 background.

### Leaf growth phenotyping

Both wild-type (Col-0) and mutant plants were grown *in vitro* on 14-cm-diameter Petri dishes at 21°C under a 16-h-day (110 mmol/(m^2^s)) and 8-h-night regime. The mutant line was grown together with the appropriate control on one plate to correct for plate effects. For each condition (MS or mannitol), 4–6 plates with 8 wild-type and 8 mutant seeds per plate were sown. Half of the plants were grown on solid (9 g/L agar, Sigma) ½ MS control medium and the other half on solid ½ MS medium with the addition of 25 mM D-mannitol (Sigma). In total, 1 and 2 independent experiments were performed for *aha2-4* and *crrsp38*, respectively. The plates were photographed at 22 DAS and all images were analysed using ImageJ (Schindelin *et al.*, 2015) to measure the projected rosette area.

## RESULTS AND DISCUSSION

### Proteome and phosphoproteome profiling to unravel the early mannitol response

To gain insight in the early molecular changes associated with mannitol-triggered osmotic response, we focused on changes in the proteome and phosphoproteome in expanding leaf tissue of *Arabidopsis thaliana*. We opted for low concentrations of the osmoticum mannitol (25 mM), which enabled a mild stress that does not affect plant survival but solely represses growth (Skirycz *et al.*, 2011*a*; Claeys *et al.*, 2014). Specifically, *A. thaliana* seedlings at 15 days after stratification (DAS) were transferred to ½ MS medium containing 25 mM mannitol and after 30 min, the third expanding leaf was harvested in 4 biological repeats. We chose the 30 min time point because this time point coincides with the earliest changes observed in the transcriptome data on mild osmotic stress in growing leaves (Van den Broeck *et al.*, 2017). Next, proteins were extracted and used for two parallel analyses: (i) the total proteome, enabling us to identify key proteins responding to mannitol, and (ii) the phosphoproteome, allowing us to gain insights into the early phosphorylation events. The proteome analysis of control and mannitol-treated samples resulted in total in the identification of 2932 protein groups (a protein group includes proteins that cannot be unambiguously identified by unique peptides but have only shared peptides) (**Figure 1** **and Supplementary Table S1**). The phosphoproteome analysis led to the identification of 3698 phosphorylated peptides that could be mapped on 1466 proteins (**Figure 1** **and Supplementary Table S2**). In this list of identified phosphopeptides, the contributions of pS, pT, and pY were 82.2%, 17.0%, and 0.8%, respectively (**Figure 1**). To address if the early mannitol-triggered leaf proteome is significantly different from a more long-term exposure, we also performed a 4 h mannitol treatment. This time point was chosen as it is 2 h later than the maximum in expression of several key transcription factors that were previously identified (Van den Broeck *et al.*, 2017). The third expanding leaf was harvested in 3 biological replicates and led to identification of 2633 proteins (**Figure 2** **and Supplementary Table S3**).

**Figure 1.**
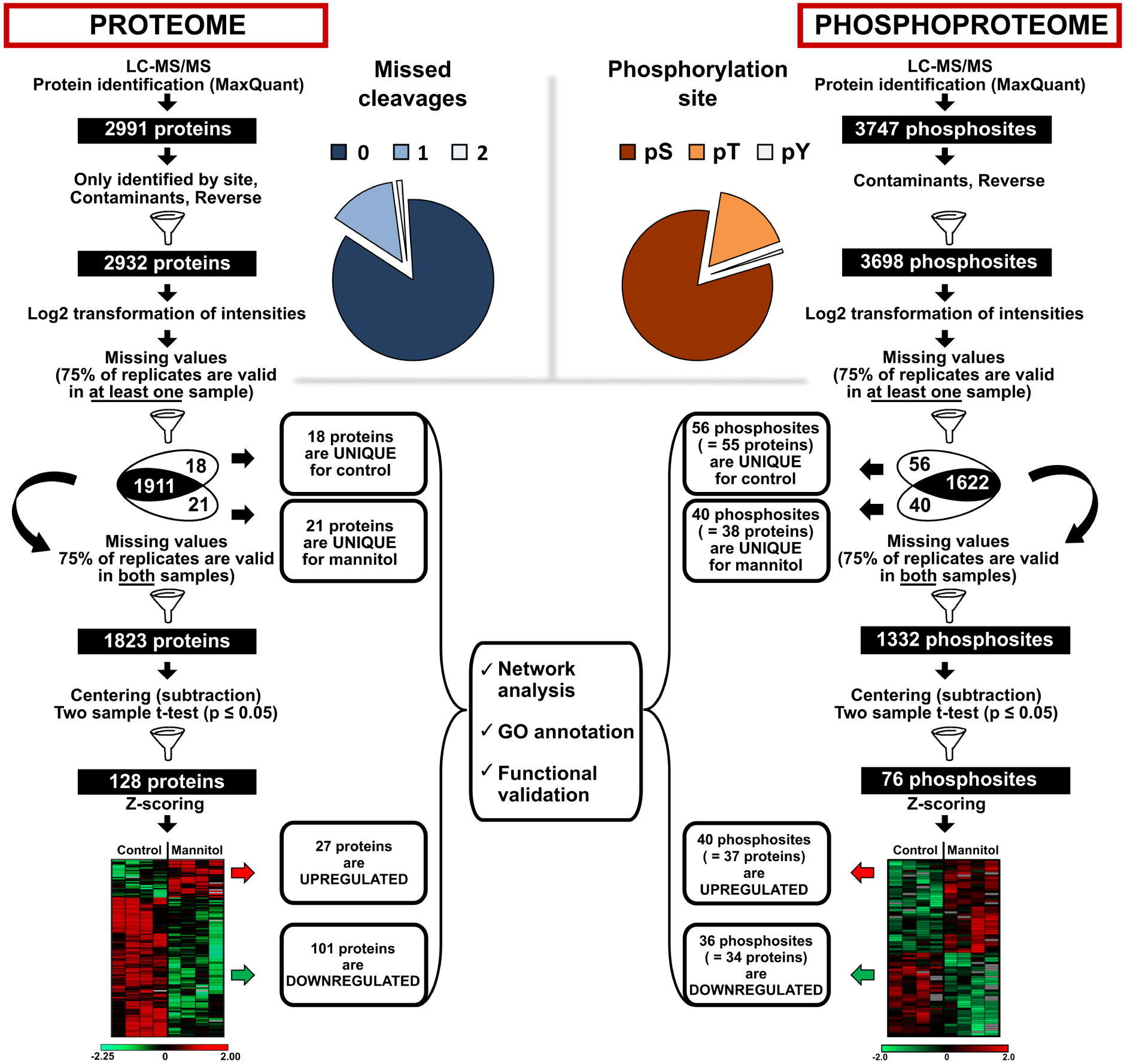
Mannitol-triggered protein and phosphoprotein changes upon short exposure (30 min) This workflow illustrates the steps to obtain a reliable set of proteins or phosphosites following LC-MS/MS. The portions for proteins and phosphosites (including the% serine (S), threonine (T) and tyrosine (Y) sites) are indicated.

**Figure 2.**
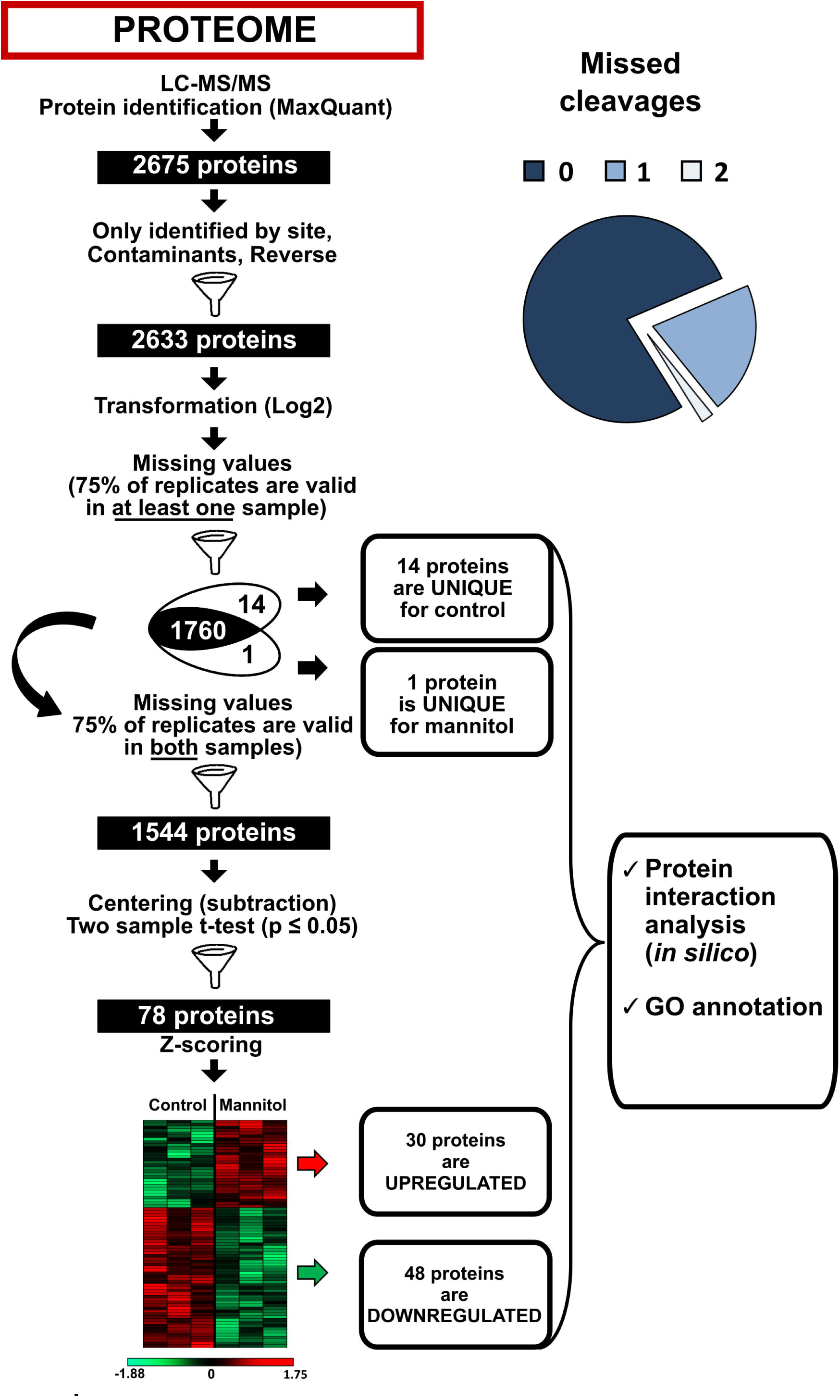
Mannitol-triggered protein changes upon long exposure (4 h) This workflow illustrates the steps to obtain a reliable set of proteins following LC-MS/MS. The numbers for detected and selected proteins are indicated.

### Data filtering approach to identify relevant candidates for further studies

One of the challenges in quantitative proteomics is dealing with missing data (Karpievitch *et al.*, 2012; Lazar *et al.*, 2016), which cannot be processed using standard regression methods (e.g., t-test and ANOVA). To overcome this, missing values can be imputed or ignored (Karpievitch *et al.*, 2012). However, this can lead to a misinterpretation of results and, more importantly, to the loss of potentially interesting proteins or phosphosites for further studies (Lazar *et al.*, 2016). A protein or a phosphorylated peptide that is present in a treated sample and absent in the control conditions (“one-state” or “unique” proteins or phosphopeptides, hereafter referred to as unique) and vice versa could be of great biological interest.

To date, post-Mass Spectrometry (MS) data analysis remains a highly debated field with numerous imputation techniques and sophisticated statistical approaches that aim to deal with a complete dataset with missing values (Koopmans *et al.*, 2014; Lazar *et al.*, 2016; Wang *et al.*, 2017*b*). Here, to overcome the problem of missing values without imputation or complex statistical analysis, we applied a hybrid approach for data analysis that treats intensity-based and presence/absence data separately (**Figures 1** **and** **2** **and Supplementary** **Figure 1**). The original, complete dataset containing *log2*-transformed intensities was split in three subsets (**Supplementary** **Figure 1; see Supplementary Information for details**). Dividing the dataset in three subsets allowed us to minimise the number of missing values in the input for regression analysis (subset 1), eliminate unreliable detections or quantifications (subset 2) and include unique proteins or phosphopeptides (subset 3) (**Figures 1** **and** **2, Supplementary** **Figure 1** **and Supplementary Tables S1-3**). The follow-up statistical workflow was performed as described previously (Vu *et al.*, 2016). In brief, the dataset was centred around zero by a subtraction of the medium within each replicate and subjected to a two-sample t-test (p < 0.05) (**Figures 1** **and** **2**).

### Early (30 min) mannitol-triggered effects on the leaf proteome

For the proteome analysis, the above-described approach resulted in 18 and 21 unique proteins detected only in the control and mannitol-treated sample, respectively (**Figure 1** **and Supplementary Table S1**). In addition, statistical analysis determined 128 differentially abundant proteins: 27 higher and 101 that were found lower in abundance upon mannitol treatment compared to the control (**Figure 1** **and Supplementary Table S1**). To simplify data characterisation further, we arranged proteins in two groups: upregulated (proteins detected only in mannitol-treated samples and significantly more abundant upon mannitol treatment) and downregulated (proteins detected only in control samples and significantly more abundant in control conditions).

Using the PLAZA 4.0 platform, GO enrichment analysis was conducted on the dataset in the context of biological process (**Supplementary Table S4**). GO enrichment on biological processes showed that differentially regulated proteins (including unique ones) were involved in various processes, such as response to stress and abiotic stimulus.

To better understand the relationships between the 167 up- and down-regulated proteins, we constructed a protein-protein interaction network consisting of 105 interacting proteins with 242 (potential) interactions (see Material and Methods for details). To get insight into the early biological processes affected by mild osmotic stress, GO annotations of up- and down-regulated proteins were superimposed on the network and nodes were grouped accordingly (**Figure 3**). This approach revealed that most interacting proteins were involved in protein metabolism, specifically in amino acid biosynthesis, translation and ribosome biogenesis, protein folding, intracellular protein transport and ubiquitin-dependent protein degradation.

**Figure 3.**
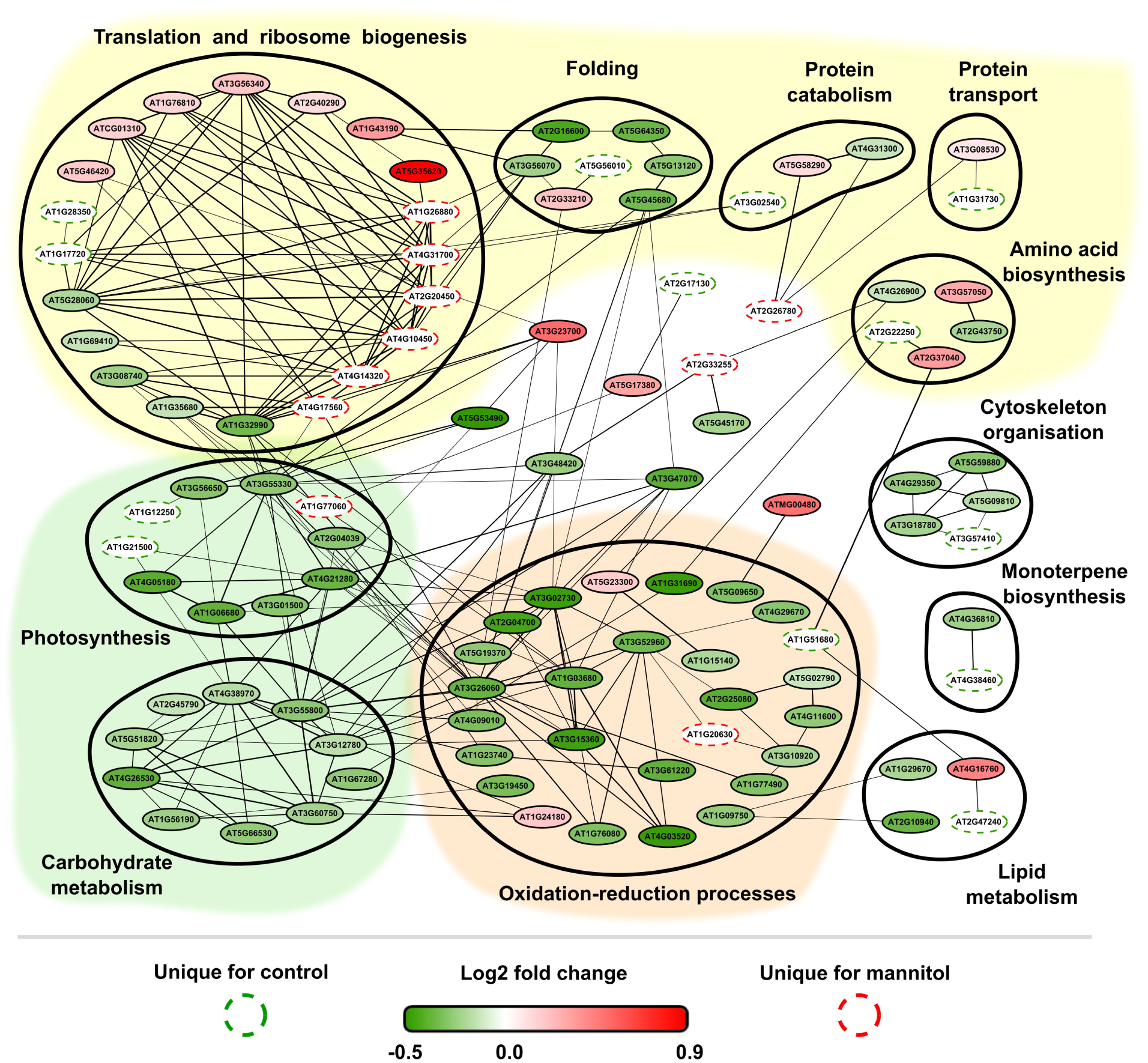
Protein-protein interaction network of significant mannitol-regulated proteins (30 min treatment). GO annotations for biological process of up- and down-regulated proteins were superimposed on the network and nodes were grouped accordingly. Colored backgrounds indicate functions related to protein metabolism (yellow), photosynthesis and carbohydrate metabolism (green) and oxidation-reduction processes (orange). Unique proteins were indicated with dashed lines while differentially abundant proteins were coloured from dark green ranging to red depending on the *log_2_* fold change. Thickness of connecting lines indicates a combined score of interaction.

In a first large group of differentially abundant interacting proteins, we observed that ribosomal proteins were highly regulated upon mannitol exposure, suggesting an altered capacity for protein translation. Six ribosomal proteins were unique for the mannitol treatment and two were upregulated. This is in line with the observation of long-term proteome changes in growing Arabidopsis leaves subjected to mannitol, where the levels of ribosomal and translational proteins were also found highly regulated, both up and downregulated (Skirycz *et al.*, 2011*b*).

A second large group of proteins in our network was involved in oxidation-reduction processes and biotic and abiotic stress responses. CAT1 was detected as a unique protein for the mannitol-treated samples, most likely protecting plant cells against toxic effects of ROS produced upon osmotic stress. Surprisingly, other reduction-oxidation protein family members (2 glutathion peroxidases, 2 ascorbate peroxidases and 9 thioredoxines, 1 periredoxin, 1 glutaredoxin and 1 dehydro-ascorbate reductase) appeared to be downregulated. This is in contrast with the long-term proteome changes in growing leaf tissue (Skirycz *et al.*, 2011*b*) and not expected because peroxidases have been widely described as ROS scavengers and are involved in the ROS damage repair (Kapoor and Sveenivasan, 1988; Caverzan *et al.*, 2012; Bela *et al.*, 2015). ROS are known to play a dual role: being toxic and destructive molecules, but, they can also serve as signalling molecules regulating stress responses, growth and development (Kovtun *et al.*, 2000; Foyer and Noctor, 2005; Vanderauwera *et al.*, 2005; Gadjev *et al.*, 2006; Pitzschke and Hirt, 2006; Brown and Griendling, 2009; Hossain *et al.*, 2015). The downregulation of these scavenger proteins might be a result of the lack of ROS production during the very early stages of mild osmotic stress or because these enzymes are a source of hydrogen peroxide (H_2_O_2_) (Blokhina *et al.*, 2003). The first hypothesis is in line with the observation that H_2_O_2_ levels do not accumulate within the first 10 min upon a hyperosmotic treatment and even decrease to levels lower than the control at later time points in Arabidopsis cell cultures (Beffagna *et al.*, 2005). On a transcript level, glutathione peroxidases have already been shown to be downregulated in rice upon drought stress (Passaia *et al.*, 2013). Deficiency in some ROS scavengers led to an increase in the plant’s sensitivity to oxidative stress while exposure to osmotic and salt stresses resulted in an increased tolerance (Miller *et al.*, 2007).

The third largest cluster in the network consisted of proteins related to photosynthesis and carbohydrate metabolism. Members of this group were mainly downregulated under mannitol stress except for a phosphoenolpyruvate carboxylase family protein (PEPC, AT1G77060). This is in agreement with previous studies where it was shown that abiotic stresses can induce *PEPC* gene expression in wheat, Arabidopsis and sorghum (Echevarría *et al.*, 2001; González *et al.*, 2003; García-Mauriño *et al.*, 2003; Sánchez *et al.*, 2006). Moreover, recent studies demonstrated that the maize *PEPC* gene was able to confer drought tolerance and increase grain yield in transgenic wheat plants (Qin *et al.*, 2016). Photosynthesis-related proteins, such as PHOTOSYSTEM II SUBUNIT Q and PHOTOSYSTEM II SUBUNIT P, were downregulated upon mannitol-induced stress, as were many chloroplast-located proteins (18 of the 121) (Lawlor and Tezara, 2009). This was expected as the photosynthetic electron transfer chain can produce ROS species (Ramachandra Reddy *et al.*, 2004).

Another interesting protein identified as unique for mannitol-treated samples was SUPER SENSITIVE TO ABA AND DROUGHT 2 (SAD2)/ ENHANCED MIRNA ACTIVITY 1 (EMA1) (**Supplementary Table S1**), an importin protein that regulates nuclear transport and is involved in the regulation of miRNA’s (Verslues *et al.*, 2006*b*; Wang *et al.*, 2011). *SAD2/EMA1* is transcriptionally not induced by ABA and stress treatments, but mutant lines show a hypersensitivity to ABA treatment and salt stress (Verslues *et al.*, 2006*b*).

To conclude, our results point towards a downregulation of photosynthesis and upregulation of CAT1 probably for ROS scavenging, both systems potentially reduce ROS production. In addition, our results suggest an unexpected new role for peroxidases in the early mannitol response as these were downregulated.

### Lack of correlation between early mannitol-triggered transcript and protein fold changes

Because some proteins are mainly regulated at post-transcriptional level, such as the above mentioned EMA1, we evaluated to what extent changes in transcript abundance for a selection of 14 genes correlated with protein levels. For this, expanding leaf tissue (leaf 3, 15 days after sowing) was harvested 20 and 40 min after mild mannitol treatment. We included the genes with the highest change in protein abundance and some of the unique proteins. Surprisingly, we found almost no obvious changes after 20 and 40 min of mannitol treatment at the transcriptional level for the genes (**Figure 4**). This suggested that changes in protein levels at 30 min were not caused by changes in transcript abundance, but more likely due to differences in protein degradation or stabilisation. We concluded that even though the transcript level was tested for only a subset of proteins, the transcriptome poorly reflects the proteome, which is in agreement with other studies (Jogaiah *et al.*, 2013; Bai *et al.*, 2015; Walley *et al.*, 2016).

**Figure 4.**
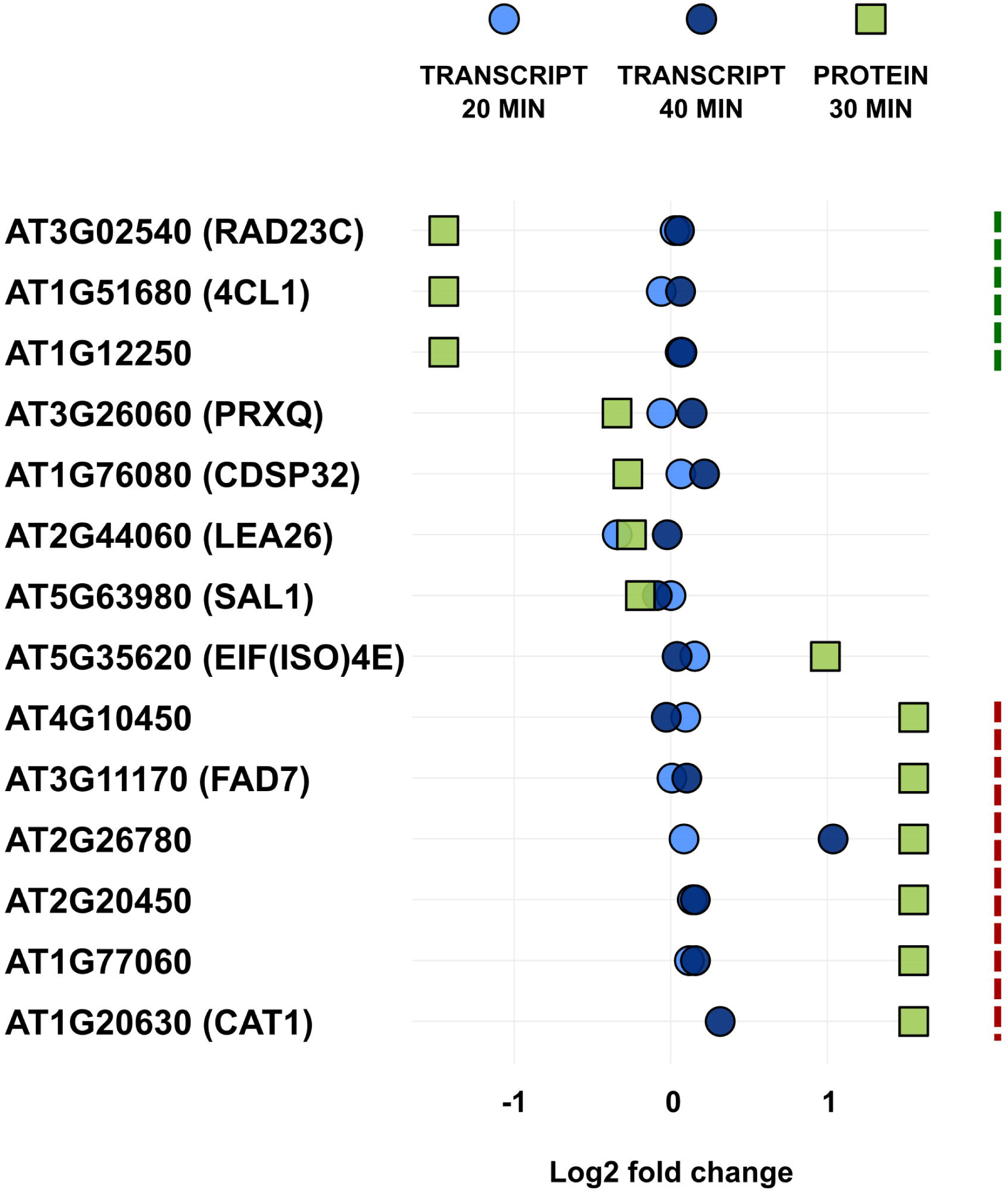
Comparison of protein abundance with transcript level of corresponding genes. The differential expression of significant differentially up- or down-regulated proteins was analysed. The expression and protein levels were measured in expanding leaf tissue upon mannitol treatment and compared to control conditions. Dashed lines indicate proteins unique for control (green) or mannitol-treated (red) samples. RAD23C – RADIATION SENSITIVE23C; 4CL1 – 4-COUMARATE:COA LIGASE 1; PRXQ – PEROXIREDOXIN Q; CDSP32 – CHLOROPLASTIC DROUGHT-INDUCED STRESS PROTEIN OF 32 KD; LEA26 – LATE EMBRYOGENESIS ABUNDANT 26; SAL1 – SAL1 phosphatase; EIF(ISO)4E – EUKARYOTIC TRANSLATION INITIATION FACTOR ISOFORM 4E; FAD7 – FATTY ACID DESATURASE 7; CAT1 – CATALASE 1.

### Effects of four hours mannitol treatment on the leaf proteome

To get an idea on changes in protein abundance upon more prolonged mild osmotic stress, we also performed a proteome analysis of growing leaf tissue after 4 hours of mannitol treatment. The above described data analysis workflow led to the identification of 15 unique proteins (based on absence or presence in 3 out of 3 biological replicates), 14 for the control and one for the mannitol-treated sample. Together with the statistical analysis, this resulted in 83 differentially abundant proteins (49 more abundant and 34 less abundant upon stress) (**Figure 2** **and Supplementary Table S3**). GO enrichment on biological processes showed that differentially regulated proteins (including unique ones) were, among several others, involved in response to stress and to abiotic stimulus (**Supplementary Table S4**). A protein-protein interaction network was built for up- and downregulated proteins (see Material and Methods for details), and similar to 30 min mannitol treatment, the vast majority of interacting proteins was involved in translation and ribosome biogenesis (**Figure 5**).

**Figure 5.**
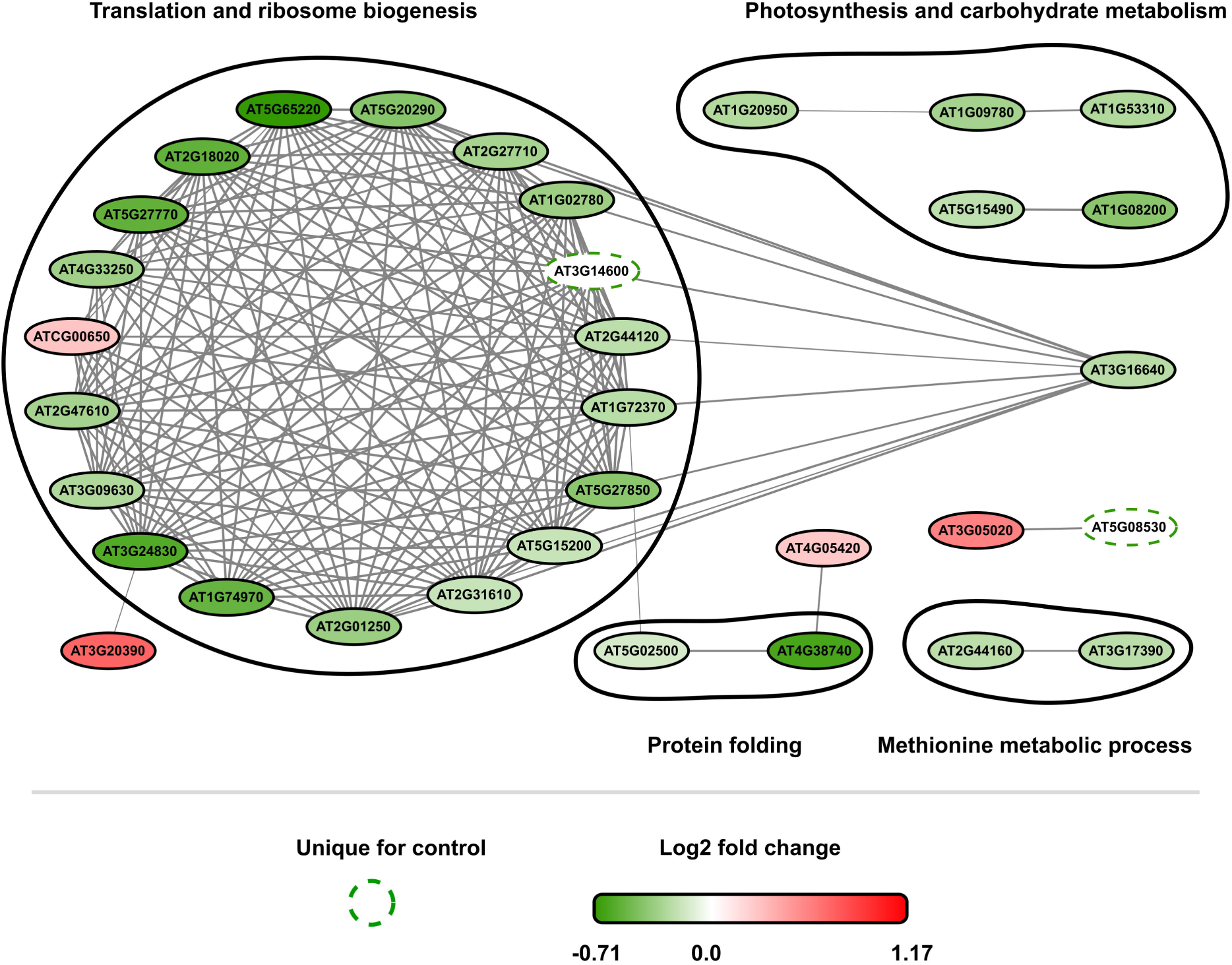
Protein-protein interaction network of significant mannitol-regulated proteins (4 h treatment). GO annotations for biological process of up- and down-regulated proteins were superimposed on the network and nodes were grouped accordingly. Unique proteins were indicated with dashed lines while differentially abundant proteins were coloured from dark green ranging to red depending on the *log_2_* fold change. Thickness of connecting lines indicates a combined score of interaction.

To unravel dynamic changes of proteins that were up- and downregulated at 30 min, we traced these proteins in the 4 h mannitol dataset. Despite that we cannot compare the exact values from both experiments, as they were processed separately, we can make some assumptions based on the protein fold changes. Thus, 167 up- and downregulated proteins of the 30 min dataset were mapped on the total 4 h proteome data and 120 proteins were retained (**Figure 6** **and Supplementary Table S5**). However, only 6 of these proteins were significantly differentially abundant after 4 h (**Supplementary Table S3**). Of the 120 mapped proteins 59 proteins showed the same trend at 30 min and 4 h after mannitol treatment of which 16 proteins remained upregulated and 43 proteins downregulated. This group of “stable” proteins contained various reduction-oxidation proteins, such as 5 thioredoxins, 2 peroxiredoxins, 1 ascorbate peroxidase, 1 glutathione peroxidase, 1 superoxide dismutase (SOD1) and 1 catalase (CAT1). Another group of proteins, consisting of 11 proteins, was upregulated at 30 minutes and downregulated after 4 h of mannitol treatment. The group of proteins that were downregulated at 30 minutes and became upregulated after 4 h of mannitol treatment included 50 proteins with diverse functions. Overall, these observations suggested a dynamic control of protein levels during mild osmotic stress response.

**Figure 6.**
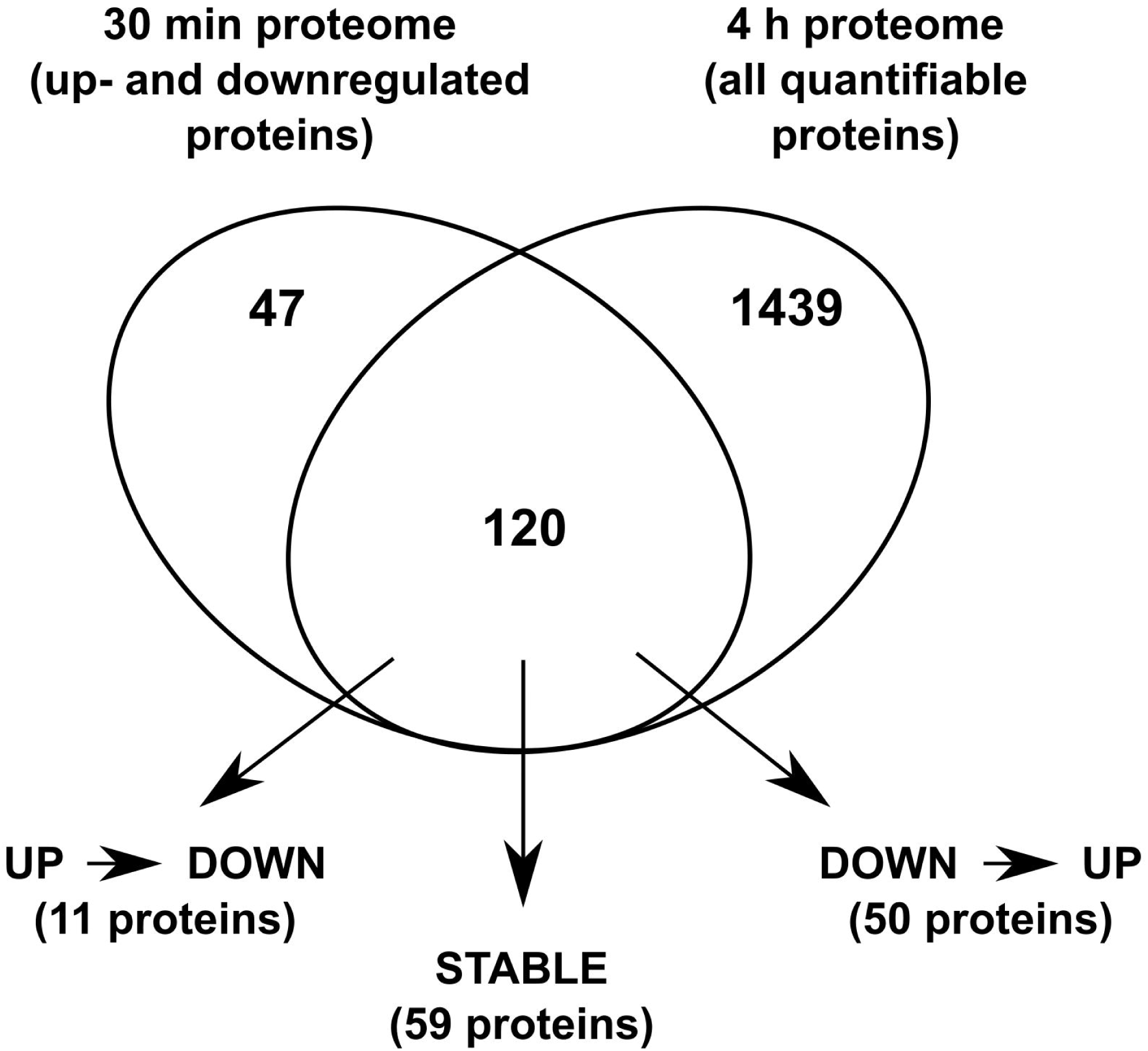
Venn diagram showing the overlap between the significant up- and down-regulated proteins from the 30 min proteome data set and all quantifiable proteins from 4 h proteome data set. In the overlap, three subsets of proteins are identified based on the changes in their abundances from 30 min to 4 h of mannitol stress; stable proteins are up- or downregulated at both 30 min and 4 h, “UP to DOWN” indicate proteins that are upregulated after 30 min but downregulated at 4 h and the “DOWN to UP” indicates the opposite.

### Early (30 min) mannitol-triggered effects on the leaf phosphoproteome

In the phosphoproteome analysis, 96 unique phosphorylated peptides were detected – 56 and 40 for control and mannitol-treated samples, respectively (**Figure 1** **and Supplementary Table S2)**. The statistical workflow resulted in a list of 76 differentially abundant phosphopeptides: 40 higher and 36 lower in abundance upon mannitol treatment compared to the control (**Figure 1** **and Supplementary Table S2**). Similar as for the proteome data, we combined unique and differentially abundant phosphopeptides in two sets of upregulated and downregulated proteins and subjected them to further analyses.

GO analysis of biological process terms revealed that proteins with differentially regulated phosphosites under mild osmotic stress (including unique phosphosites) were involved in several processes, including (protein) phosphorylation, regulation of cellular response to stress, response to abscisic acid and mannitol metabolic process (**Supplementary Table S4.**).

To unravel possible interactions between the proteins that were differentially phosphorylated and unique, we constructed, as for the proteome datasets, a protein-protein interaction network consisting of 44 interacting phosphoproteins with 42 (potential) interactions combined with GO categorisation for biological processes (see Material and Methods for details). This approach revealed that interacting phosphoproteins were mainly involved in photosynthesis, splicing, chromatin remodelling and transport (**Figure 7**). In general, this network analysis showed that 27% of the detected phosphoproteins are part of a subnetwork, suggesting that upon a stimulus, more than one protein of the subnetwork is differentially phosphorylated, potentially affecting its activity. However, it should be noted that we cannot pinpoint a role for the identified change in phospho-status in relation to the protein function or activity.

**Figure 7.**
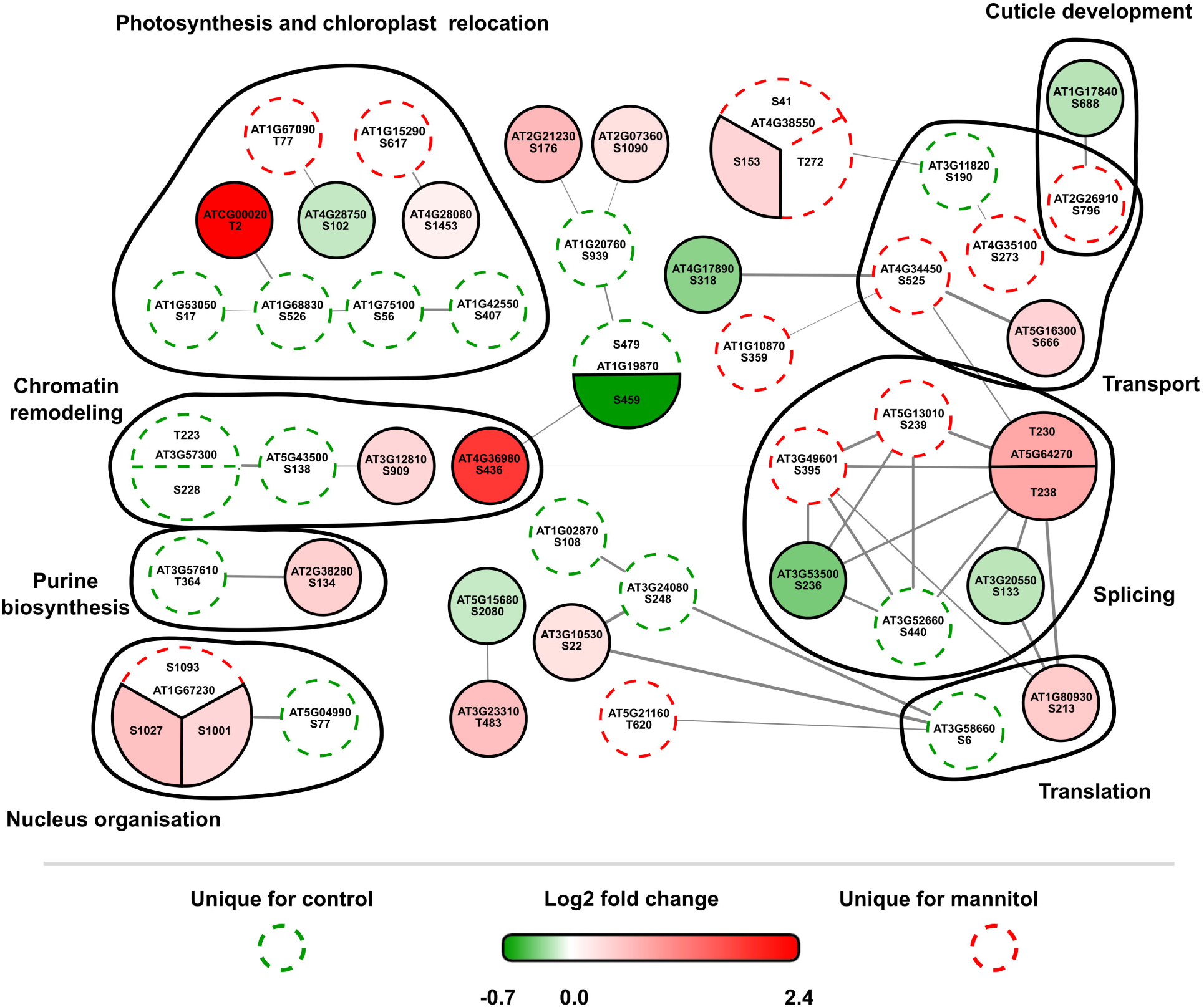
Protein-protein interaction network of significant mannitol-regulated phosphopeptides mapped on the corresponding proteins (30 min treatment). GO annotations for biological process of up- and down-regulated proteins were superimposed on the network and nodes were grouped accordingly. Unique phosphosites were indicated with dashed lines while differentially abundant phosphosites were coloured from dark green ranging to red depending on the *log_2_* fold change. Thickness of connecting lines indicates a combined score of interaction.

Upon mild osmotic stress, the DELLA proteins play an important role in the growth inhibition in leaves upon mild osmotic stress (Claeys *et al.*, 2012). GA binds to its receptor GA INSENSITIVE DWARF1 (GID1) (Ueguchi-Tanaka *et al.*, 2005) and forms a complex with DELLA proteins, leading to its degradation (Nakajima *et al.*, 2006). Under mild stress, the gene encoding a GA-degradation enzyme, *GA2-OX6*, is upregulated by several mannitol responsive transcription factors (Van den Broeck *et al.*, 2017), which leads to a decrease in GA levels and subsequently to the stabilisation of DELLA proteins. Such stabilisation of DELLA proteins leads to changes in the activity of transcription factors and expression of GA-regulated genes. In our phosphoproteomic dataset, we found a few differentially phosphorylated proteins that are linked to DELLAs. For example, bZIP16 (AT2G35530; Ser^152^, >1.5-fold downregulated) (**Table 1**), a transcriptional repressor, has been shown to directly repress *REPRESSOR OF GA-LIKE 2* (*RGL2*), encoding a DELLA protein (Hsieh *et al.*, 2012). We thus hypothesize that a decrease in phosphorylation of bZIP16 might decrease its activity and its repression on *RGL2*. Another example is the CALCIUM-DEPENDENT PROTEIN KINASE (CDPK/CPK)-RELATED PROTEIN KINASE 2 (TAGK2/CRK2) (AT3G19100), for which a phospho-site at Ser^57^ was upregulated 1.6-fold upon mannitol treatment (**Table 1**). Recently it was shown that TAGK2/CRK2 phosphorylates the GA RECEPTOR RING E3 UBIQUITIN LIGASE (GARU) at Tyr^321^which results in the disruption of the interaction between GARU and the GA RECEPTOR GA INSENSTIVE DWARF1 (GID1). Because this interaction is disrupted, GARU is not able to induce the degradation of GID1. *TAGK2*-overexpressing plants thus show an increased GID1 stabilisation and DELLA degradation (Nemoto *et al.*, 2017). However, the role of phosphorylation associated with TAGK2 has not yet been studied.

**Table 1.**
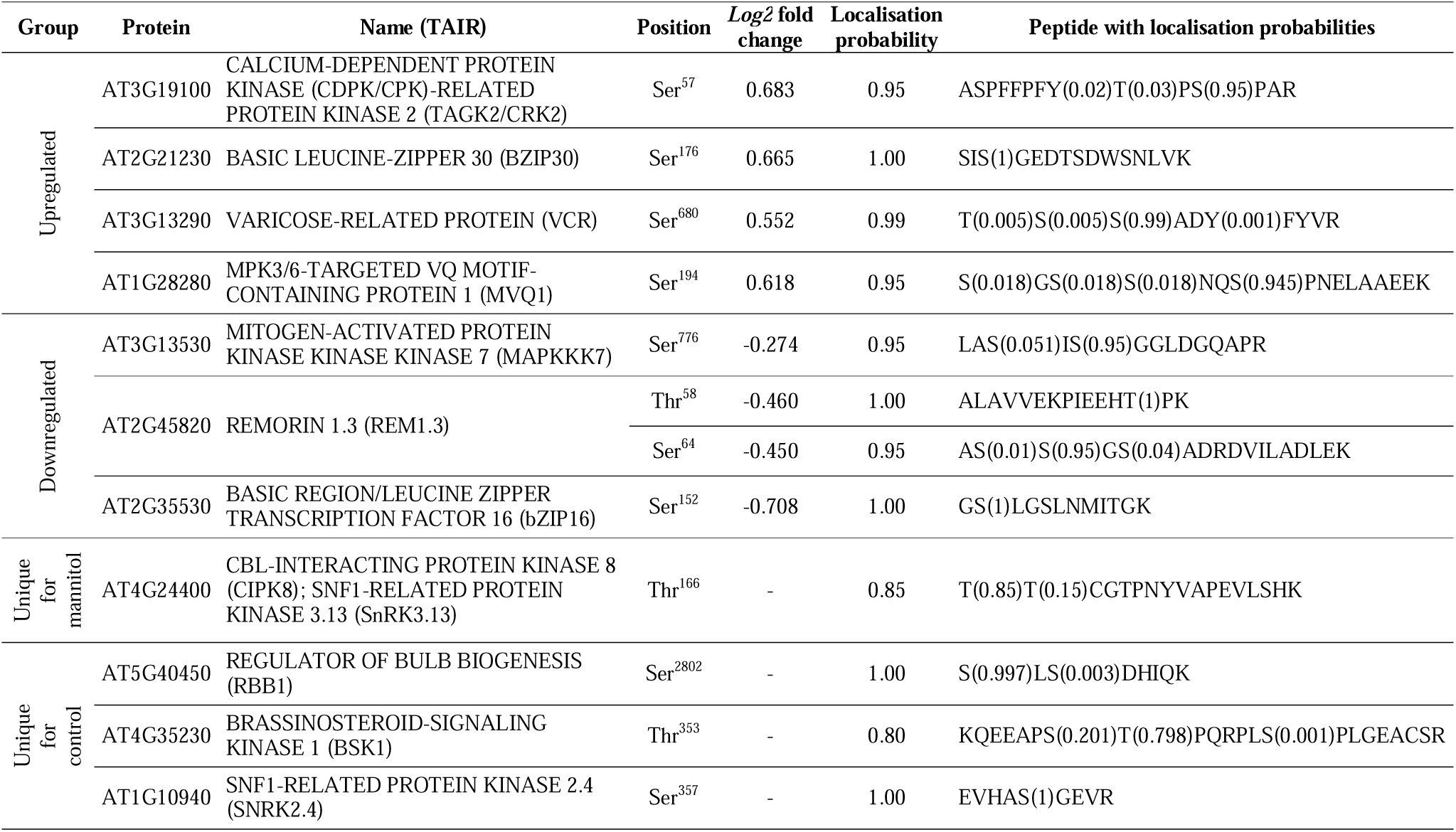
Phosphorylated sites mentioned in text.

Considering the interest in early signalling cascades underlying responses to mild osmotic stress, we focused on proteins with up- and downregulated phosphosites possessing protein kinase activity. Using the HMMER online tool (Finn *et al.*, 2015), we found 7 and 10 (potential) protein kinases of which phosphosites were up- and downregulated, respectively (**Supplementary Table S6**). Upregulation of phosphorylation sites of a kinase can indicate an activation of the kinase itself and, as a consequence, its downstream signaling cascades; and dephosphorylation can imply the opposite (Wang *et al.*, 2007; Tarrant and Cole, 2009; Day *et al.*, 2016). Next, we looked into predicted overrepresented kinase motifs as the sequence consensus of phosphopeptide motifs reflects the kinase-specific regulation of substrate phosphorylation and the identity of the corresponding kinases (**Supplementary Figure S2**). This revealed that for the upregulated phosphopetides, the protein motifs [SP] and [TP] were the most enriched motifs. Peptides containing the proline (P)-directed [SP] and [TP] motifs are suggested to be targets of MAP-kinases, SnRK2, RLKs, CDPKs, CDKs, AGC family protein kinases and STE20-like kinases (van Wijk *et al.*, 2014). For the downregulated phosphopeptides, [SP] and [RxxS] motifs were overrepresented, of which the [RxxS] motif is recognized by MAP kinases.

Some examples of the identified kinases that have a changed phospho-status upon mild mannitol stress are CALCINEURIN B-LIKE PROTEIN (CBL) – INTERACTING PROTEIN KINASE 8 (CIPK8, AT4G24400), MAP kinase kinase kinase 7 (MKKK7, AT3G13530), brassinosteroid-signalling kinase 1 (BSK1, AT4G35230) and SnRK2.4 (AT1G10940) (**Table 1**). CIPKs are involved in calcium signalling cascades and are found to be induced in response to stress (Hu *et al.*, 2009; Pandey *et al.*, 2014). Mitogen activated protein kinase (MAPK) cascades are known to be involved in drought stress (Ichimura *et al.*, 2000) and in PAMP signalling (Asai *et al.*, 2002; Pitzschke *et al.*, 2009). The detected MKKK7 was demonstrated as a negative regulator of flg22-triggered signalling and basal immunity (Mithoe *et al.*, 2016), but could have other functions as well. BSK1 is a target of the brassinosteroid receptor BRASSINOSTEROID INSENSITIVE 1 (BRI1) and plays a role in the brassinosteroid signalling during plant immunity (Shi *et al.*, 2013). SnRK2.4 is a well-known regulator of the salt stress response in plants. According to recent findings, SnRK2.4 belongs to the SnRK2 group 1 of which its members are not activated by abscisic acid (ABA) (Kulik *et al.*, 2011). The importance of phosphorylation for SnRK2.4 activity was previously demonstrated upon salt stress and the abolishment of SnRK2.4 activity in mutant lines led to an increased sensitivity to salt (Krzywińska *et al.*, 2016). The first phase of salt stress is osmotic stress, strengthening a role for SnRK2.4 in this process (Shavrukov, 2013). However, our analysis showed a different mannitol-regulated phosphorylation site than those previously described (Kline *et al.*, 2010), namely Ser^357^. The Ser^357^ residue was dephosphorylated upon mannitol treatment, and is located in a protein-protein interaction motif outside the activation loop of the kinase that has not been linked to osmotic stress responses (Kulik *et al.*, 2011).

### A normalised early mannitol-triggered differential leaf phosphoproteome

A common question related to quantitative phosphoproteomics is whether the measured phosphorylation changes result from changes in kinase or phosphatase activity or from changes in phosphoprotein abundance. In the context of quantitative phosphoproteomics it is important to correct – to the extent possible – the measured phosphorylation changes with respect to changes in protein abundance (Vu *et al.*, 2016). Taking into account that phosphopeptide abundance is directly depending on protein abundance, we normalised phosphosite intensities. It should be noted that this analysis is only possible for a subset of identified phosphosites because as a result of the enrichment for phosphorylated proteins during the phosphoproteome analysis, most of them are not detected in the whole proteome where no enrichment was performed. Furthermore, it should be mentioned that this normalised phosphoproteome data set does not imply that the non-normalised phosphopeptides are not interesting to investigate; merely that these changes could not be corrected for protein abundance. 172 up- and downregulated phosphorylated peptides derived from 158 phosphoproteins were mapped on the total proteome data and 32 proteins were found to be overlapping. Next, the *log_2_* fold change of the protein was subtracted from the *log_2_* fold change of the phosphosite. This defined a set of phosphorylation events that are fully due to changes in kinase and phosphatase activity and not due to changes in protein abundance (to the extent these changes in phosphorylation status do not impact on protein abundance) (**Figure 8** **and Supplementary Table S7**). Sixteen phosphosites could not be normalised as they belonged to a group of unique phosphosites and did not have a fold change value. Within the normalised phosphopeptides, two phosphosites of a remorin family member REM1.3 (AT2G45820), Thr^58^ and Ser^64^, were down regulated upon mannitol treatment (**Figure 8** **and Table 1**). REM1.3 was already reported to be phosphorylated upon oligo-galacturonide treatment which elucidates a plant stress response (Kohorn *et al.*, 2016). REM1.3 is proposed as a scaffolding protein for signalling at the plasma membrane and is thus an interesting candidate for mannitol-induced signalling (Marín *et al.*, 2012).

**Figure 8.**
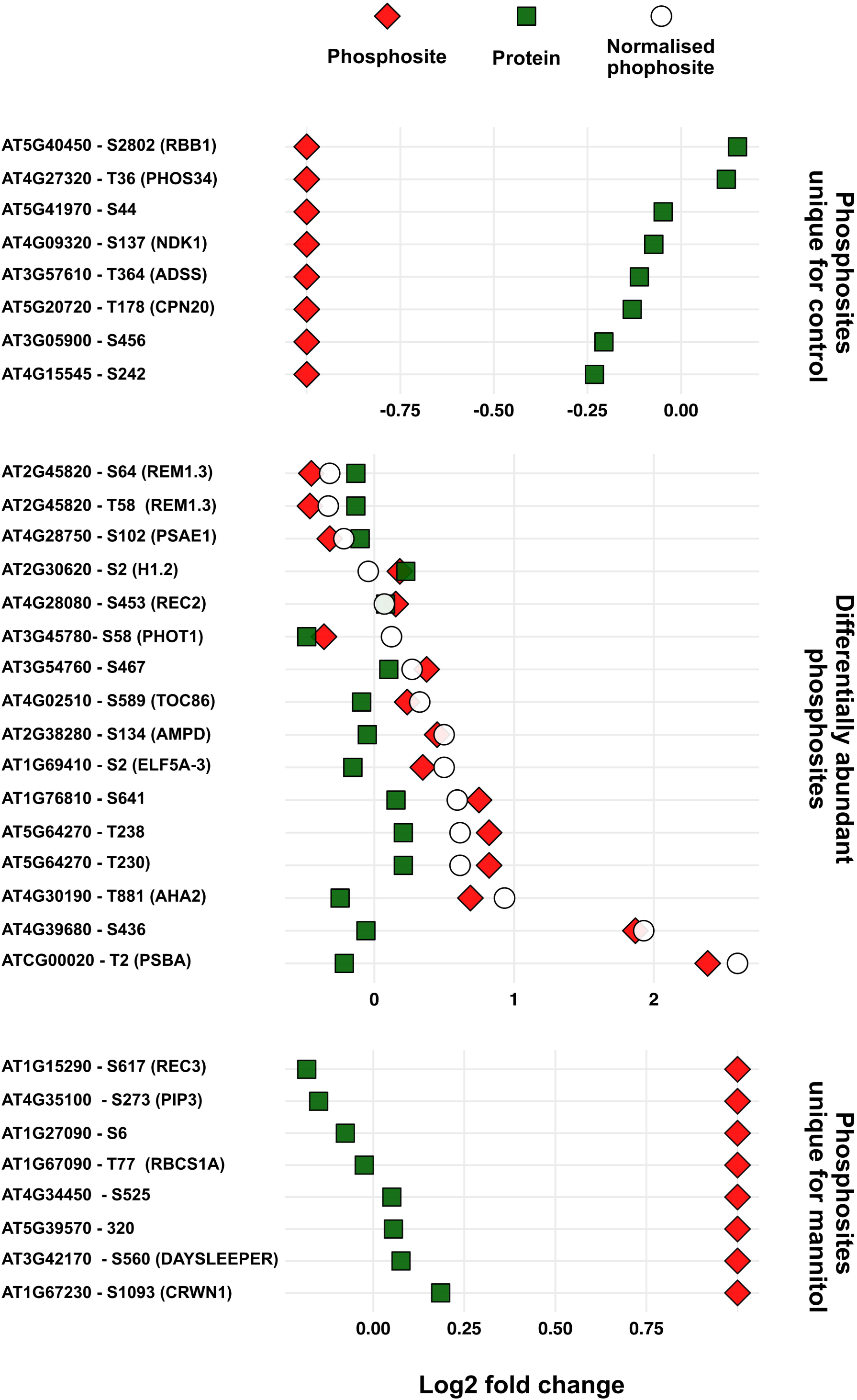
A normalised mannitol-triggered phosphoproteome. Significantly up- and downregulated phosphosites were normalised by subtracting the *log_2_* fold change of the protein abundance from the *log_2_* fold change of the phosphosite, with the exception of the unique phosphosites. In total, 32 differentially phosphorylated proteins could be mapped on total proteome data. RBB1-REGULATOR OF BULB BIOGENESIS1; PHOS34 – Phosphorylated protein of 34 kDa; NDK1 – NUCLEOSIDE DIPHOSPHATE KINASE 1; ADSS – ADENYLOSUCCINATE SYNTHASE; CPN20 – CHAPERONIN 20; REM1.3 – Remorin 1.3; PSAE1 – Photosystem I reaction center subunit IV A; H1.2 – HISTONE 1.2; REC2 – REDUCED CHLOROPLAST COVERAGE 2; PHOT1 – PHOTOTROPIN 1; TOC86 – TRANSLOCON AT THE OUTER ENVELOPE MEMBRANE OF CHLOROPLASTS 86; AMPD – ADENOSINE 5’-MONOPHOSPHATE DEAMINASE; ELF5A-3 – EUKARYOTIC ELONGATION FACTOR 5A-3; AHA2 – H(+)-ATPASE 2; PSBA – PHOTOSYSTEM II REACTION CENTER PROTEIN A; REC3 – REDUCED CHLOROPLAST COVERAGE 3; PIP3 – PLASMA MEMBRANE INTRINSIC PROTEIN 3; RBCS1A – RIBULOSE BISPHOSPHATE CARBOXYLASE SMALL CHAIN 1A; DAYSLEEPER – Zinc finger BED domain-containing protein; CRWN1 – CROWDED NUCLEI 1.

### Comparative phosphoproteome analysis identifies bZIP30 and RBB1 as general players in osmotic stress response

To assess whether the mannitol-regulated phosphorylated proteins in growing leaf tissue are part of a more general stress response or are specific for the stress-induced growth-regulating response, we compared our data set (30 min 25 mM mannitol) with 3 previously published osmotic stress-related phosphoproteome datasets (Xue *et al.*, 2013; Stecker *et al.*, 2014; Bhaskara *et al.*, 2017*b*). Given the differences in experimental set-up (long-term versus short-term or mild versus severe stress) (see **Table 2** for details), we found little overlap between all four datasets; only two proteins, BASIC LEUCINE-ZIPPER 30 (BZIP30)/DRINK ME (DKM) and REGULATOR OF BULB BIOGENESIS1 (RBB1) (Han *et al.*, 2015), were detected in all datasets, possibly indicating a general role in osmotic stress responses (**Figure 9** **and Supplementary Table S8**). Interestingly, the same phosphorylation site, Ser^176^, of bZIP30 was found to be upregulated in the 3 datasets where mannitol was used. It should be noted that bZIP30 (Ser^176^, >1.5-fold upregulated, **Table 1**) was also identified in our network as a central phosphoprotein in the longest interaction chain (**Figure 7**). This transcription factor influences the expression of cell cycle and cell expansion genes, two processes that are affected by mild mannitol treatment (Skirycz *et al.*, 2011*b*; Lozano-Sotomayor *et al.*, 2016). In addition, we found 4 common mannitol-responsive phosphoproteins (based on overlap between (Xue *et al.*, 2013), (Stecker *et al.*, 2014) and this study). The phosphorylation site Ser^680^ of VARICOSE-RELATED PROTEIN (VCR, AT3G13290) was identified as upregulated in all studies where mannitol was used. In addition, six phosphoproteins were exclusively found in the two mild osmotic stress studies (Bhaskara *et al.*, 2017*a* and this study), including a mitogen activated protein kinase kinase kinase-like protein (AT3G58640) and MPK3/6-TARGETED VQ MOTIF-CONTAINING PROTEIN 1 (MVQ1, AT1G28280).

**Table 2.** Details on phosphoproteomic studies for comparative analysis

| Study | Material | Growth medium | Compound | Concentration | Time | Setup | Method |
| --- | --- | --- | --- | --- | --- | --- | --- |
| Current study | Leaf #3 of 15-days-old seedlings | Agar plates | Mannitol | 25 mM | 30 min | Transfer | Label-free |
| Bhaskara et al., 2017 | 7-day-old seedlings | Agar plates | PEG | a low water potential (21.2 MPa) | 96 h | Transfer | ITRAQ |
| Stecker et al., 2014 | 10-day-old seedlings | Liquid culture | Mannitol | 300mM | 5 min | Medium replacement | <sup>15</sup>N-labelling |
| Xue et al., 2013 | 12-day-old seedlings | Liquid culture | Mannitol | 800mM | 30 min | Transfer | Label-free |

**Figure 9.**
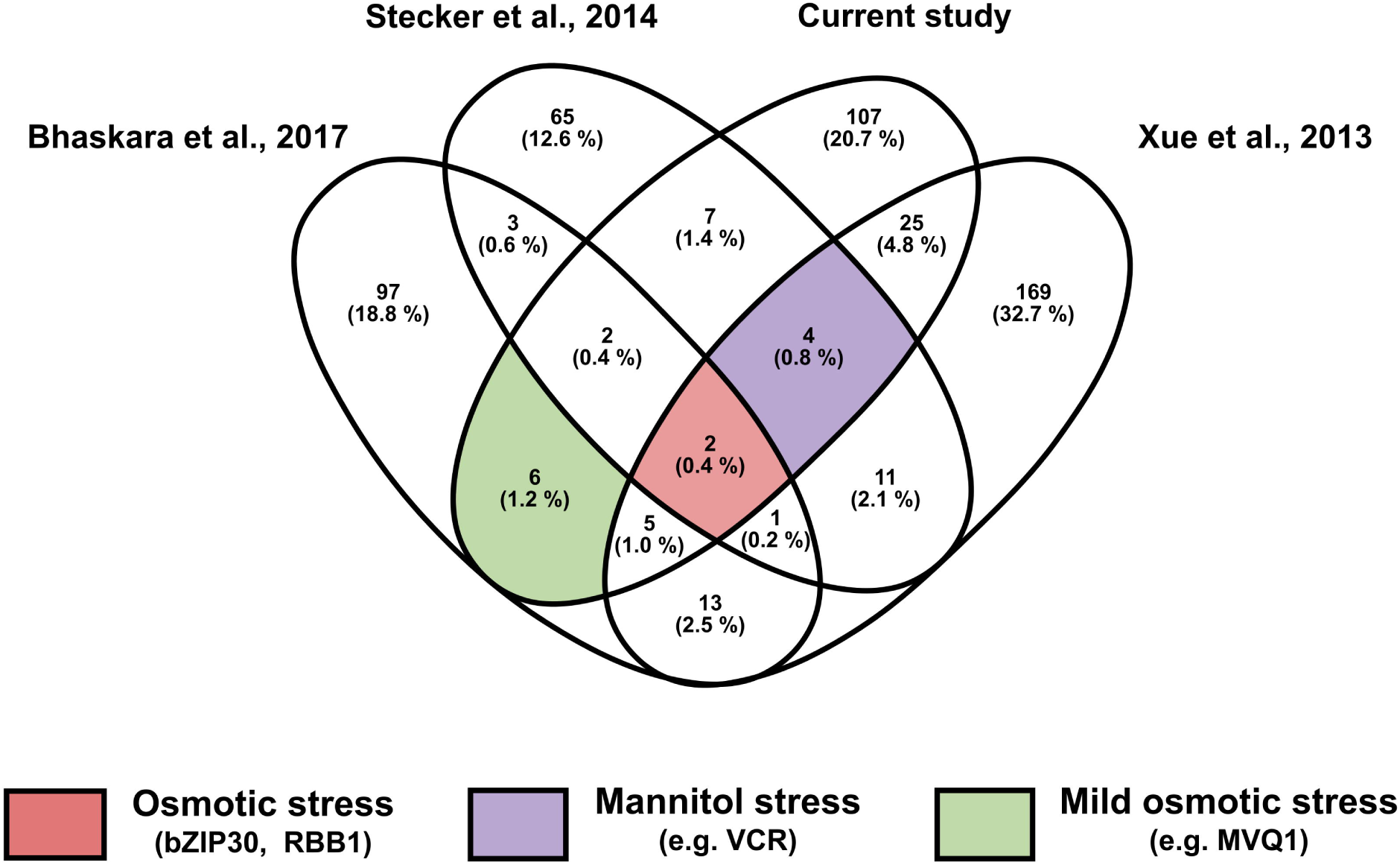
Venn diagram showing the overlapping phosphoproteins from four recent phosphoproteomic studies of osmotic stress responses, including our current study. Details on selected studies, concerning the osmoticum and concentration that was used, are indicated in Table 2. bZIP30 – BASIC LEUCINE-ZIPPER 30; RBB1-REGULATOR OF BULB BIOGENESIS1; VCR – VARICOSE-RELATED PROTEIN; MVQ1 – MPK3/6-TARGETED VQ MOTIF-CONTAINING PROTEIN 1.

Apart from the different methodologies, the limited overlap points out the large difference between short and long-term phosphorylation response upon stress, indicating the transient and dynamic behaviour of phosphorylation events. Additionally, the severity of the stress determines the phosphorylation response (Bhaskara *et al.*, 2017*a*).

### (Phospho) proteome profiling identifies AHA2 and CRRSP38 as growth regulators under mannitol stress

To assess the involvement of proteins with a differential abundance or phosphorylation status in shoot growth and/or mannitol response, and thus the quality of our data set, we selected two candidates for phenotypic analysis. From the mannitol-regulated proteome data set (**Supplementary Table S1**), we selected the uncharacterized CYSTEINE-RICH REPEAT SECRETORY PROTEIN 38 (CRRSP38, AT3G22060). Interestingly this protein was identified as down-regulated after 30 min of mannitol application and was not detected after 4 h of mannitol stress treatment. This is in contrast to the absence of any mannitol-induced transcriptional change at 20 and 40 min, and the increasing expression from 2 h on (**Supplementary Figure S4**). Possibly, this is due to a transcript – protein level feedback mechanism.We obtained a knock-out allele of CRRSP38 (SALK_151902), referred to as *crrsp38-1* (**Figure 10** **and Supplementary Figure S3**). Phenotypic analyses of *crrsp38-1* at 22 days after sowing revealed a larger rosette area both under control and mild mannitol conditions (**Figure 10**), suggesting that this protein is potentially a general growth regulator instead of specifically involved in the growth regulation upon mild stress. It will be interesting to explore the molecular function of CRRSP38 in leaf growth regulation, and its precise role upon mild mannitol-induced stress.

**Figure 10.**
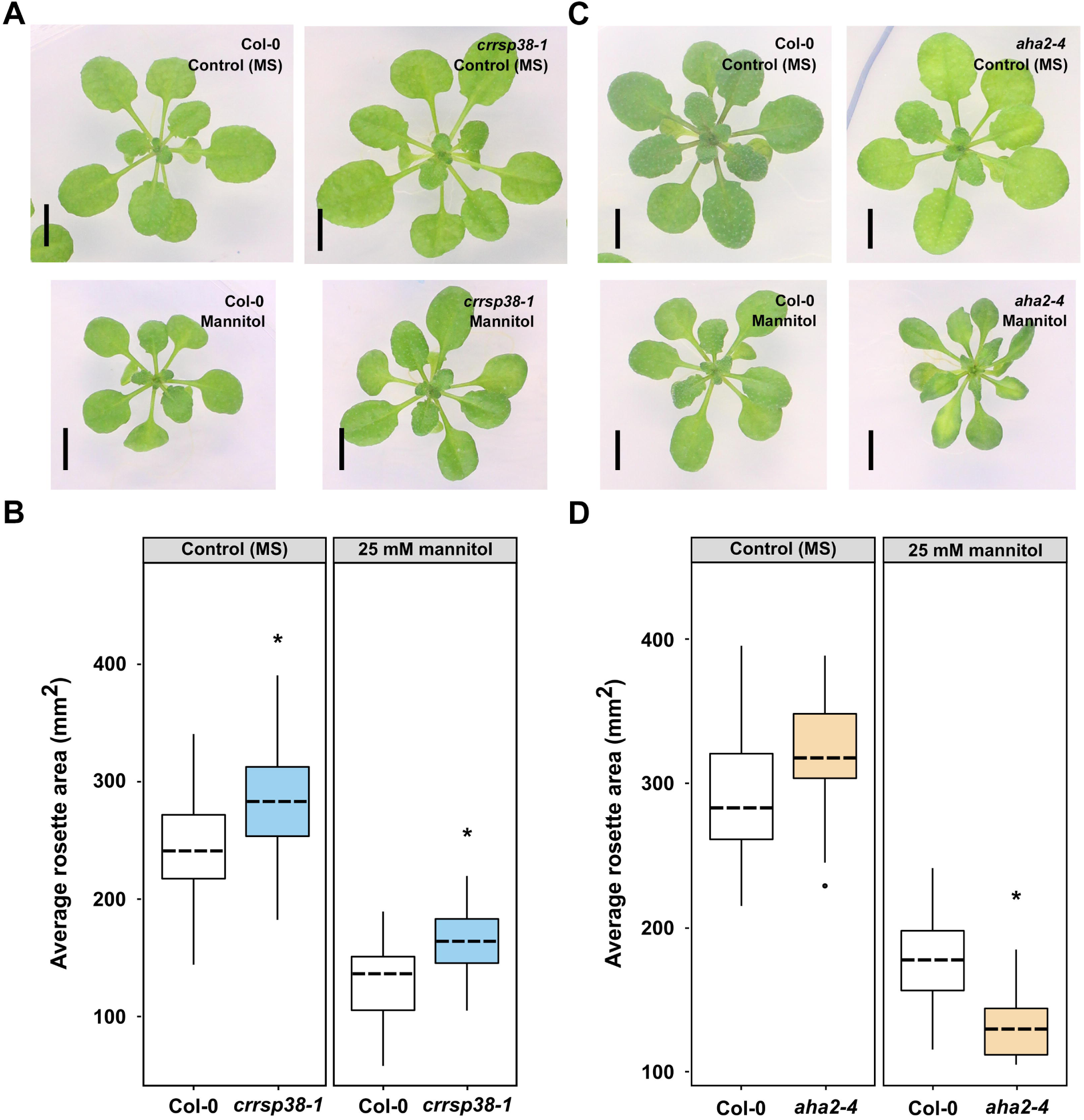
Phenotypes of T-DNA insertion lines for selected candidates. **(A, C)** Representative pictures of *crrsp38-*1 and *aha2-4* lines compared to Col-0 at 22 days after stratification grown on control (MS) or mannitol (25 mM). Scale bar, 5 mm. **(B, D)** Quantification of the rosette area of *crrsp38-1* (B) and *aha2-4* (D) at 22 days after stratification grown on control (MS) or 25 mM mannitol (MA). Boxplots are combined values of at least 30 seedlings from 4–6 different plates and 2 or 1 independent experiments for *crrsp38-1* or *aha2-4*, respectively. Asterisk indicates a significant difference with p < 0.05 based on ANOVA. In addition, the ANOVA analysis indicated that the genotype:treatment interaction for *aha2-4* was significant (p < 0.05).

From the normalised phosphoproteome, we focused on AHA2 (AT4G30190), an H(+)-ATPASE 2 of which phosphosite Thr^881^ showed an 1.9-fold increase in phosphorylation after mannitol treatment and had not yet been connected to mannitol stress, and does not show any obvious mannitol-induced transcriptional change in a time course experiment (**Figure 8**, **Supplementary Figure S4 and Supplementary Table S7**). Thr^881^ is situated in the conserved autoinhibitory region I of the C-terminal domain of AHA2 (Rudashevskaya *et al.*, 2012; Falhof *et al.*, 2016), and rapid PLANT PEPTIDE CONTAINING SULFATED TYROSINE 1 (PSY1)-induced *in planta* phosphorylation of AHA2 at Thr^881^ increases proton efflux (Fuglsang *et al.*, 2014). We hypothesized AHA2 might play a role in growth inhibition upon mild osmotic stress. Therefore, we characterized the strong knock-down *aha2-4* mutant with only about 10% *AHA2* expression compared to wild type (Haruta *et al.*, 2010). As for *crrsp38-1*, the rosette area of the *aha2-4* mutant was measured at 22 DAS under normal and mild mannitol conditions. The *aha2-4* plants have a slightly larger rosette area in control conditions (8%) and were significantly more sensitive to mannitol compared to wild-type (**Figure 10**). Specifically, the *aha2-4* mutant showed a significant reduction of 59% under mannitol conditions compared to a reduction of 38% in the wild-type plants. This suggested that AHA2 can indeed regulate the mannitol-induced growth inhibition.

In summary, our (phospho)proteome-centred approach allowed the identification of novel mannitol stress-related players.

## CONCLUSIONS

In this study, label-free proteomic and phosphoproteomic analyses were performed on expanding Arabidopsis leaves exposed to mild osmotic stress. By performing the proteome analysis at two different time points, 30 min and 4 h, dynamic patterns in protein abundance could be observed. In general, after 30 min of stress ribosomal proteins were upregulated upon mannitol treatment, and photosynthesis and reduction-oxidation-related proteins were downregulated. While after 4 h, ribosomal proteins were downregulated. Furthermore, the lack of correlation between transcriptional changes prior to changes in protein abundance points towards an important role of protein degradation/stabilisation upon stress. In addition, we identified several proteins that had an altered phosphorylation status upon mild osmotic stress, suggesting an important role for kinase and phosphatase-mediated signalling. We also identified several important regulators, such as the transcription factor bZIP30, which is likely a central component of both mild and severe osmotic stress. However, previously and in this study, no transcriptional changes were observed for bZIP30, indicating that by solely studying transcriptomics, central proteins involved in stress response are likely missed. On the other hand, several transcription factors from the recently described mild mannitol stress-associated gene regulatory network were not identified in our (phospho)proteomes. In addition, we identified proteins that were specifically phosphorylated under short-term mild osmotic stress, such as AHA2. Phenotypic analysis of an *aha2* knock-out mutant indeed confirmed a role for AHA2 in the regulation of growth upon mild osmotic stress.

Taken together, our datasets further stress the importance of proteome-and phosphoproteome-based approaches, in addition to transcriptomics, for unravelling the molecular mechanisms underlying growth regulation under stress.

## SUPPLEMENTARY DATA

### Supplementary Information

**Supplementary Figure S1.** Visual explanation for the 3 subsets described in the main text.

**Supplementary Figure S2.** Visual depiction of predicted overrepresented kinase motifs based on Motif X analysis.

**Supplementary Figure S3.** Details of *crrsp38-1* T-DNA line.

**Supplementary Table S1.** Proteins identified at 30 min after mannitol treatment, including raw data, differentially abundant and unique proteins.

**Supplementary Table S2.** Phosphosites identified at 30 min after mannitol treatment, including raw data, differentially abundant and unique phosphosites.

**Supplementary Table S3.** Proteins identified at 4 h after mannitol treatment, including raw data, differentially abundant and unique proteins.

**Supplementary Table S4.** GO enrichment on biological processes.

**Supplementary Table S5.** Comparative analysis of proteome datasets from 30 min (significantly different and unique proteins) and 4 h (all dataset) mannitol treatment.

**Supplementary Table S6.** Protein kinases with differentially phosphorelated and unique phosphosites predicted by HMMER.

**Supplementary Table S7.** Phosphorylated proteins normalised for protein abundance.

**Supplementary Table S8.** Comparative analysis of outputs from 4 phosphoproteomic studies on osmotic stress.

## ACKNOWLEDGEMENTS

We thank Lam Dai Vu and Veronique Storme for valuable discussions. L.V.d.B. is a predoctoral fellow of the Research Foundation Flanders (FWO no. 131013). This research received funding from the Bijzonder Onderzoeksfonds Methusalem Project (BOF08/01M00408).

